# Reduced activity of Adenylyl Cyclase 1 Attenuates Morphine Induced Hyperalgesia and Inflammatory Pain in Mice

**DOI:** 10.1101/2020.12.02.408419

**Authors:** Kayla Johnson, Alexis Doucette, Alexis Edwards, Val J. Watts, Amanda H. Klein

## Abstract

Opioid tolerance and opioid-induced hyperalgesia during repeated opioid administration and chronic pain are associated with upregulation of adenylyl cyclase activity. The objective of this study was to test the hypothesis that a reduction in adenylyl cyclase 1 (AC1) activity or expression would attenuate morphine tolerance and hypersensitivity, and inflammatory pain using murine models. To investigate opioid tolerance and opioid-induced hyperalgesia, mice were subjected to twice daily treatments of saline or morphine using either a static (15 mg/kg, 5 days) or an escalating tolerance paradigm (10-40 mg/kg, 4 days). Systemic treatment with an AC1 inhibitor, ST03437 (5 mg/kg, ip), reduced morphine tolerance and morphine hyperalgesia in mice. Lumbar intrathecal administration of a vector incorporating adeno-associated virus and short-hairpin RNA against *Adcy1* reduced morphine induced hypersensitivity compared to control vector treated mice. In contrast, morphine antinociception, along with baseline thermal paw withdrawal latencies, motor performance, exploration in an open field test, and burrowing behaviors were not affected by intrathecal *Adcy1* knockdown. Knockdown of *Adcy1* by intrathecal injection also attenuated inflammatory mechanical hyperalgesia after intraplantar administration of Complete Freund’s Adjuvant (CFA) after one week post injection. This *Adcy1* knockdown strategy also increased burrowing and nesting activity after CFA injection when compared to controls. Together, these data indicate targeting AC1 to mitigate opioid-induced adverse effects, or as a method to treat chronic pain, are appropriate as a clinical approach and further development into generating pharmaceuticals targeting these genes/proteins may prove beneficial in the future.

## Introduction

Opioids are one of the most common analgesics used to alleviate pain clinically by inhibiting neuronal signal transmission through the mu opioid receptor (MOR). Individuals with chronic pain use opioids on a daily basis for pain management, causing the development of analgesic tolerance, leading to dosage escalation. In the clinic, tolerance is defined as a requirement for increased opioid doses to maintain analgesia. Opioid-induced hypersensitivity is defined as the increased sensitivity to pain as a result of chronic opioid use. Whether the increased opioid requirement is caused by the decreasing analgesic efficacy of the drug, as in tolerance, or by an increase in spontaneous pain or lowering the nociceptive threshold, the clinical effect is the same [19]. Furthermore, if the patient ceases therapy, there is a possibility for withdrawal and nerve hypersensitivity, increasing the likelihood for opioid dependence and abuse situations.

Upon agonist binding to the MOR, adenylyl cyclase (AC) is inhibited thereby blocking the formation of cyclic adenosine monophosphate (cAMP). However, prolonged agonist stimulation of the MOR leads to an inability to inhibit AC, or a phenomenon called heterologous sensitization of AC, causing intracellular activity of AC to increase, thereby increasing intracellular levels of cAMP [43]. Enhancement of cAMP levels due to prolonged opioid exposure has long been connected to opioid tolerance and opioid dependence in both *in vitro* [7; 33] and *in vivo* studies, particularly in the spinal cord and dorsal root ganglia (DRG)[9; 29]. More recently, the activation of Ca^2+^/calmodulin ACs, particularly AC1 and AC8, were implicated in the initial stages of morphine tolerance and withdrawal as AC1 and AC8 knockout mice have increased latencies during the first few days of morphine tolerance testing as well as decreased withdrawal behaviors [25; 47]. AC1 and AC8 have also been linked to the development of both acute and chronic persistent inflammatory pain [16; 17; 36; 42] and a global loss of either AC1 or AC8, or in combination, appear to have a role in attenuating morphine tolerance and withdrawal [37; 40]. However in an inflammatory pain model in mice, loss of AC1, but not AC8, decreased nocifensive responses to formalin[36]. AC isoform-selective pharmacological inhibitors have been developed, particularly for AC1, and appear to attenuate chronic pain in mice [4; 21; 26; 39]. Of note ST034307, a highly selective inhibitor for AC1, has shown effectiveness at providing analgesia in a mouse model of inflammatory pain [4]. To date, it is unknown if selectively inhibiting AC1 activity or reducing AC1 expression after chronic MOR stimulation alters the development of opioid tolerance and opioid-induced hypersensitivity.

The purpose of this study was to better understand the activity of AC1 during morphine tolerance, opioid induced hypersensitivity and chronic inflammatory pain. To accomplish this, pharmacological inhibition of AC1 or a short hairpin RNA (shRNA) knockdown strategy using adeno associated virus (AAV9) vector was used to decrease activity/expression of *Adcy1*. Systemic treatment with ST03437, reduced morphine tolerance and morphine hyperalgesia in mice. Similarly, behavioral measures indicate intrathecal administration of a viral vector expressing *Adcy1* shRNA in mice attenuates morphine tolerance, opioid-induced hypersensitivity, and decreases evoked pain measures in a mouse model of inflammatory pain.

## Materials and Methods

### Animals

All experimental procedures involving animals were approved and performed in accordance with the University of Minnesota Institutional Animal Care and Use Committee guidelines. Adult male C57Bl6 mice were obtained via Charles River (5-6 weeks old, Raleigh, NC). Mice were acclimated to individual testing apparatuses prior to behavioral testing. Mice were euthanized by isoflurane anesthesia (5%) followed by decapitation at the end of the study.

### Tissue Collection and mRNA Isolation

Tissues harvested from animals were flash frozen in liquid nitrogen and stored at −80 °C. Total mRNA was isolated from tissues using Tri Reagent (T9424, Sigma Aldrich, St. Louis, MO) and RNeasy Mini Kit (Qiagen, Germantown, MD) according to manufacturer’s protocol with DNase digestion. Complimentary DNA synthesis was performed with 50 ng total mRNA using Omniscript RT Kit (Qiagen, Germantown, MD) and random nonamers (Integrated DNA Technologies, Coralville, Iowa) according to manufacturer’s protocol.

### Quantitative PCR

Quantitative PCR was performed using SYBR Green I dye with LightCycler 480 technology (Roche, Branchburg, NJ, USA). The cDNA copy number was typically quantified against a ≥5 point, 10-fold serial dilution of a gene specific cDNA standard. Internal controls included negative RT-PCR samples and comparative expression versus a housekeeping gene, *18S*. Fold expression of each gene of interest was determined by: (mean gene concentration/mean 18s concentration)/(mean gene concentration in saline/mean 18s concentration in saline). See Supplemental Table 1 for gene specific primers used.

### Drugs and Delivery

Morphine (Sigma Chemical, St. Louis, MO) was administered through a 100 uL subcutaneous injection in saline. ST034307 (6271, Tocris Bioscience, Minneapolis, MN) was dissolved in 10% β-cyclodextrin with 5% DMSO in saline and administered 5 mg/kg through a 100 uL intraperitoneal injection 15 minutes after morphine administration. Morphine efficacy was determined using an escalating dose response curve (5-20 mg/kg) waiting 30 minutes after each injection [22]. For morphine tolerance experiments, baseline mechanical paw withdrawal testing was performed before administration of 15 mg/kg morphine for five days [27]. Escalating morphine tolerance was performed similarly, except increasing doses of morphine, starting at 10 mg/kg and increasing 10mg/kg/day, were administered over the course of four days. In each model, morphine was delivered twice per day (~0800 and ~1800 hours). Mechanical threshold testing was performed 30 and 60 minutes post morphine administration in the morning. To determine the degree of opioid-induced hyperalgesia, paw withdrawal thresholds were assessed starting ~18 hours after the last dose of morphine. Complete Freund’s Adjuvant (CFA, F5881, Sigma Chemical, St. Louis, MO) was administered through an intraplantar injection (20 uL, undiluted) into the left hind paw. [23; 24].

### Mechanical Paw Withdrawal

Mice were acclimated to testing environment on at least two separate occasion for 30 to 60 minutes before formal testing. Testing environment consisted of a mesh floor, allowing access to animal hind paws, and individual clear acrylic chambers. Mechanical paw withdrawal (MPW) thresholds were determined by use of electronic von Frey testing equipment (Electric von Frey Anesthesiometer, 2390, Almemo® 2450, IITC Life Science, Woodland Hills, CA). The plantar surface of the hind paws were gently pressed with the probe until a nocifensive response (i.e. paw lifting, jumping, and licking) was elicted. Baseline measurements (in grams) were collected five times from both the right and left hind paw and averaged, with an interstimulus interval of at least one minute.

### Thermal Paw Withdrawal

Testing environment consisted of a glass floor heated to 30°C with individual clear acrylic chambers. The modified Hargreaves method was used to measure thermal paw withdrawal (TPW) latency (Plantar Test Analgesia Meter, 400, IITC, Woodland Hills, CA) [5]. TPW latencies were determined by the amount of time (in seconds) a heat radiant beam of light focused on the plantar surface of the hind paw was required to elicit a nocifensive response (e.g. paw lifting, shaking, and licking). A maximum time limit of 20 seconds exposure to the beam was used to avoid tissue damage. Baseline measurements were collected five times from both the right and left hind paw and averaged, with an interstimulus interval of at least two minutes.

### Adeno-Associated Virus Serotype 9 (AAV9)-Mediated Adcy1 Knockdown

Gene knockdown of *Adcy1* using shRNA was achieved using AAV9-GFP-U6-m-*Adcy1*-shRNA with AAV9-GFP-U6-scramble-shRNA as control viral vector (shAAV-251792 and 7045, titer: 1.4×10^13^ GC/mL, in PBS with 5% glycerol, Vector Biolabs, Malvern, PA, United States). Vectors were delivered by direct lumbar puncture (10 uL) in awake mice and behavioral assessments were performed 4–8 weeks post injection [13; 38].

### Rotarod Performance Test

Agility assessment was conducted using Rotamex-5 automated rotarod system (0254-2002L, 3cm rod, Columbus Instruments, Columbus, OH). Mice were placed onto a stationary knurled PCV rod suspended in the air. The initial rotation speed of 4 rpm was gradually increased by 1 rpm in 30-second intervals until animals fell off the rod or reached a speed of 14 rpm (300 seconds). Two tests were administered per animal and averaged.

### Burrowing Testing

Mice were acclimated to empty burrowing tubes for ~2 hours on at least two separate occasions before formal testing. The burrows were made from a 6 cm diameter plastic pipe and 5 cm machine screws were used to elevate the open end by 3 cm [11]. During testing, each mouse was placed in an individual cage with a burrowing tube containing 500 g of pea gravel. The amount of gravel remaining in the tube after 2 hours was used to calculate the total percent of gravel displaced from the burrow.

### Open Field Testing

The open field testing arena consisted of a 40 × 40 cm box with a white floor and black walls. Animals were placed in the open field arena, in a room with controlled adjustable lighting, and baseline activity was recorded for 30 minutes (Sony Handycam, HDR-CX405, Sony Corp., Tokyo, Japan). The distance traveled, time spent immobile, average velocity, and the change in orientation angle were computed by using data output from the Ethowatcher computational tool software (Laboratory of Bioengineering of the Institute of Biomedical Engineering and the Laboratory of Comparative Neurophysiology of the Federal University of Santa Catarina, UFSC, available: http://ethowatcher.paginas.ufsc.br/) [10].

### Nesting

Mice were individually housed in clean plastic cages containing cob bedding with food and water *ad libitum* overnight. A single 2” Nestlets™ (Ancare Corp., Bellmore, NY) square was weighed and added to each cage. The next morning (~14 hours) untorn pieces of each nesting square were weighed and the resulting nests were photographed and scored on a 5-point scale as described previously [11]. Briefly the scoring system was: 1 = >90% intact, 2 = partially torn, 3 = mostly shredded but no identifiable nest, 4 = >90% torn but flat nest site, 5 = >90% torn with resulting crater nest. Scores with 0.5 units were used for nests with scores in between the aforementioned intervals.

### Microscopy

Histological sections were taken in spinal cord, DRG and sciatic nerves in order to verify the delivery of the AAV9 vector within the lumbar intrathecal space. Verification of virus inoculation were visible by the presence of green fluorescent protein (GFP). Sections (10 uM, Leica CM3050) were mounted onto electrostatically charged slides and images were collected using a Nikon TiS Microscope and associated software.

### C-fiber Compound Action Potentials

Compound action potentials (CAPs) were measured from both left and right desheathed sciatic nerves from AAV9-GFP-U6-m-*Adcy1*-shRNA and AAV9-GFP-U6-scramble-shRNA 8 weeks after intrathecal injection. Sciatic nerves were dissected from the hind limbs of mice and recordings were performed the day of harvesting. Each nerve was mounted in a chamber filled with superficial interstitial fluid composed of 107.7 mM NaCl, 3.5 mM KCl, 0.69 mM MgSO_4_, 26.2 mM NaCO_3,_ 1.67 mM NaH_2_PO_4_, 1.5 mM CaCl_2_, 9.64 mM Na^+^ gluconate, 5.5 mM d-glucose, and 7.6 mM sucrose, pH 7.4 (bubbled with 95% O_2_, 5% CO_2_). Electrical stimulation was performed at a frequency of 0.3 Hz with electric pulses of 100-μs duration at 100-10,000 uA delivered by a pulse stimulator (2100, AM Systems, Carlsborg, WA). Evoked CAPs were recorded with electrodes placed ~5 mm from the stimulating electrodes. Dapsys software was used for data capture and analysis (Brian Turnquist, Bethel University, St. Paul, MN, *www.dapsys.net)*. The stimulus with the lowest voltage producing a detectable response in the nerve was determined the threshold stimulus. The stimulus voltage where the amplitude of the response no longer increased was determined to be the peak amplitude. The conduction velocity was calculated by dividing the latency period, the time from stimulus application to neuronal initial response, by the stimulus-to-recording electrode distance.

### Data analysis

Data were collected by personnel blinded to the animal condition and treatment. The appropriate t-test, one-way, two-way, or repeated measures ANOVA followed by Bonferroni’s *post hoc* analysis was used to determine significance for MPW thresholds and TPW latencies, gene expression, burrowing, open field testing, rotarod assessments, and CAP recordings. Nonparametric tests were used for nesting behaviors. All statistical analyses were carried out using GraphPad Prism versions 7.0 and 8.0 (GraphPad Software, San Diego, CA). All other data is presented as mean ± SEM with p < 0.05 considered statistically significant.

## Results

### Adcy1 mRNA expression is increased in in the peripheral nervous system and spinal cord in mice after chronic administration of morphine

Chronic exposure of the MOR to agonists causes a decreased inhibitory response by the MOR and increases the adenylyl cyclase/cyclic-AMP activity[44]. We attempted to confirm these findings by analyzing the expression of *Adcy1* mRNA in nervous system tissues of morphine tolerant mice. The mRNA expression of adenylyl cyclase isoforms and other downstream intracellular targets were analyzed in both central and peripheral nervous system tissues including brain stem (BS), trigeminal ganglion (TG), spinal cord (SC), dorsal root ganglion (DRG) and sciatic nerve (SN) after chronic morphine treatment using qRT-PCR. To induce morphine tolerance, morphine was administered twice daily (15 mg/kg, sc in saline) for five days. The overall fold change in gene expression was calculated for morphine tolerant mouse tissues compared to control saline treated mice (Supplemental Figure 1). An increased expression of *Adcy1* is seen in SN, DRG, and TG, and a >2-fold increase was found in in the SC (Table 1). A >2-fold increase in mRNA expression was also seen in the SN for *Adcy3*, *Adcy6* and *Rapgef3* (Table 1). This data suggests AC1 may play a role in morphine tolerance in both the central and peripheral nervous systems. To further understand the physiological role of AC1 in tolerance and withdrawal during chronic morphine administration, pharmacological and gene knock-down strategies were implemented with behavioral assays.

**Table 1.** Altered levels of AC isoforms and downstream targets during morphine tolerance. Morphine tolerance was induced in mice by twice daily injections of 15 mg/kg morphine in saline (100 μL, subcutaneous) for five days. Expression of each gene in sciatic nerve (SN), dorsal root ganglion (DRG), spinal cord (SC), brainstem (BS) and trigeminal ganglion (TG) was analyzed using qRT-PCR. The mean gene concentration within each tissue was first normalized to *18S* before being compared to the same tissue from saline treated mice resulting in overall fold change. Median fold change and the range of fold change values are reported (n=4/group).

| Gene | Tissue | Fold Change | Range | Gene | Tissue | Fold change | Range |
| --- | --- | --- | --- | --- | --- | --- | --- |
| *Adcy1* | SN | 1.52 | 0.91 to 3.45 | *Adcy8* | SN | 1.68 | 0.68 to 2.64 |
|  | DRG | 1.19 | 0.83 to 2.66 |  | DRG | 1.31 | 1.12 to 1.60 |
|  | SC | 2.15 | 1.45 to 3.39 |  | SC | 0.67 | 0.47 to 0.71 |
|  | BS | 0.35 | 0.11 to 0.45 |  | BS | 0.44 | 0.23 to 0.56 |
|  | TG | 1.15 | 1.10 to 1.42 |  | TG | 1.71 | 1.16 to 2.33 |
| *Adcy2* | SN | 0.71 | 0.60 to 1.80 | *Prkaca* | SN | 1.53 | 1.13 to 2.10 |
|  | DRG | 1.56 | 1.27 to 1.68 |  | DRG | 1.52 | 1.30 to 1.91 |
|  | SC | 1.04 | 0.71 to 1.30 |  | SC | 0.91 | 0.65 to 0.98 |
|  | BS | 0.54 | 0.32 to 0.65 |  | BS | 0.5 | 0.34 to 0.61 |
|  | TG | 0.71 | 0.85 to 2.13 |  | TG | 1.08 | 0.99 to 1.13 |
| *Adcy3* | SN | 2.22 | 1.42 to 3.42 | *Prkacb* | SN | 1.42 | 1.14 to 1.90 |
|  | DRG | 1.45 | 1.14 to 1.66 |  | DRG | 1.4 | 1.32 to 1.52 |
|  | SC | 0.54 | 0.36 to 0.63 |  | SC | 0.68 | 0.50 to 0.81 |
|  | BS | 0.37 | 0.13 to 0.48 |  | BS | 0.53 | 0.33 to 0.66 |
|  | TG | 1.11 | 1.01 to 1.35 |  | TG | 1.17 | 0.99 to 1.23 |
| *Adcy5* | SN | 1.81 | 0.83 to 2.20 | *Rapgef3* | SN | 2.34 | 1.16 to 3.52 |
|  | DRG | 0.92 | 0.79 to 1.13 |  | DRG | 1.25 | 1.07 to 1.76 |
|  | SC | 0.94 | 0.68 to 1.14 |  | SC | 1.21 | 0.73 to 1.48 |
|  | BS | 0.58 | 0.37 to 0.77 |  | BS | 0.56 | 0.42 to 0.68 |
|  | TG | 1.29 | 1.07 to 1.36 |  | TG | 1.46 | 1.37 to 1.78 |
| *Adcy6* | SN | 2.13 | 1.09 to 3.16 | *Rapgef4* | SN | 1.04 | 0.80 to 1.83 |
|  | DRG | 1.35 | 0.70 to 2.25 |  | DRG | 1.47 | 1.22 to 1.56 |
|  | SC | 0.09 | 0.03 to 0.12 |  | SC | 0.9 | 0.75 to 1.19 |
|  | BS | 0.64 | 0.46 to 1.00 |  | BS | 0.8 | 0.75 to 0.83 |
|  | TG | 0.93 | 0.71 to 1.40 |  | TG | 1.07 | 0.91 to 1.28 |

### Systemic ST034307 administration attenuates morphine tolerance and withdrawal

Previous research demonstrated ST034307 acts as an AC1 inhibitor and as an analgesic in a mouse chronic inflammatory pain model [4]. The data presented here also confirm the possible antinociceptive properties of ST034307. In both mechanical and thermal nociceptive tests, the peak threshold and latency measurements increased after intraperitoneal administration of ST034307 (Figure 1A, B). A significant difference in TPW latency was seen between vehicle and ST034307 over time (Figure 1B; repeated measures ANOVA with Bonferroni’s *post hoc* test, F (1, 12) = 15.06, p = 0.0022, CI_15min_ = 1.282 to 5.912). This data demonstrates the peak antinociceptive action of ST034307 occurring around ~15 minutes post injection, but this effect is fairly weak in naïve mice. So, any changes in thresholds seen during morphine tolerance testing should be due to AC1 inhibition and not analgesia caused by ST034307 administration.

**Figure 1.**
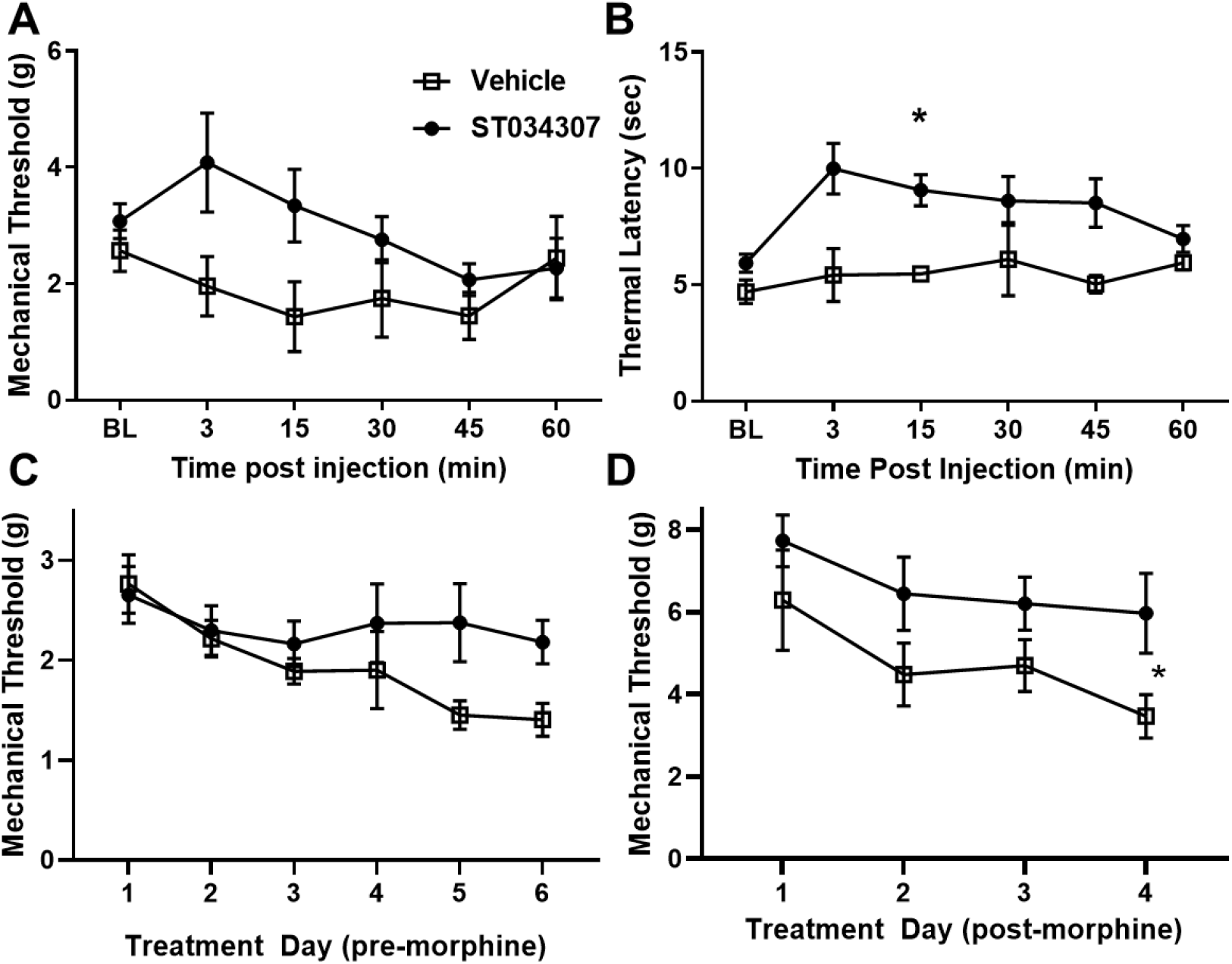
ST034307 Produces Mechanical and Thermal Antinociception and Attenuates Morphine Tolerance. To obtain mechanical thresholds and thermal paw withdrawal latencies, mice were given intraperitoneal injections of vehicle (□) or ST034307 (●) following baseline measurements (BL). (A) No significant analgesic differences were seen between vehicle and ST034307 treated mice. (B) A significant difference in thermal paw withdrawal was seen between vehicle and ST034307 treated mice (repeated measures ANOVA with Bonferroni’s *post hoc* test, F (1, 12) = 15.06, p = 0.0022, CI_15min_ = 1.282 to 5.912). To induce morphine tolerance, mice received twice daily injections of morphine (10 mg/kg on day 1 increasing to 40 mg/kg by day 4, subcutaneous, 100 uL) along with an injection of either vehicle or ST034307 (5 mg/kg, intraperitoneal, 100 uL) 15 minutes post-morphine. BL measurements were measured every morning before morphine injection (C) and 30 minutes post injection (D) with day 5 and day 6 thresholds measured ~18 hours and ~42 hours, respectfully, after last morphine injection. MPW thresholds pre-morphine were not significantly different between the two treatment groups (C; two-way ANOVA with Bonferroni’s *post hoc* test, F (1, 16) = 3.940, p = 0.0646) but mice given ST034307 had significantly higher MPW thresholds compared to vehicle treated post-morphine (D; two-way ANOVA Bonferroni’s *post hoc* test, F (1, 16) = 6.512, p = 0.0213). Asterisk indicates statistical significance (p < 0.05). Data presented as mean ± SEM with an n=4-10/group.

Previous research demonstrated ST034307 acts as an AC1 specific inhibitor capable of blocking heterologous sensitization of AC1 after chronic MOR activation *in vitro*[4]. To determine if ST034307 attenuates morphine tolerance and opioid-induced hypersensitivity *in vivo*, mice were subjected to twice daily morphine injections (10 mg/kg on day one increasing 10 mg/kg each day to a final concentration of 40 mg/kg, sc) in combination with either an injection of vehicle or ST034307 (5 mg/kg, ip), 15 minutes post-morphine. MPW thresholds were measured before the start of injections (Figure 1C) and 30 minutes post morphine injection (Figure 1D) to measure opioid-induced hypersensitivity and morphine tolerance, respectively. Although the administration of ST034307 increased paw withdrawal thresholds after morphine administration had ceased after Day 4, no significant difference is seen between the two treatment groups pre-morphine (Figure 1C; two-way ANOVA with Bonferroni’s *post hoc* test, F (1, 16) = 3.940, p = 0.0646). Mice treated with ST034307 demonstrated significantly higher MPW thresholds after morphine injections compared to vehicle treated mice (Figure 1D; two-way ANOVA Bonferroni’s *post hoc* test, F (1, 16) = 6.512, p = 0.0213), indicating that the pharmacological inhibition of AC1 can aid in the attenuation of tolerance.

### Intrathecal knockdown of Adcy1 attenuates morphine tolerance and opioid-induced hypersensitivity

An AAV9 viral vector strategy utilizing shRNA targeting to Adcy1 was used to reduce *Adcy1* expression within the peripheral nervous system and spinal cord via intrathecal injection. To ensure the shRNA knockdown strategy of the AAV9-*Adcy1* viral vector was successful, animals were sacrificed eight weeks post viral vector injections and SC and DRG tissues were collected. The mRNA copy numbers of *Adcy1* were significantly reduced in AAV9-*Adcy1* viral vector injected mice in both the SC (Figure 2A, unpaired t-test, p = 0.0201) and DRG (Figure 2A, unpaired t-test, p = 0.0370) compared to mice treated with the AAV9-scramble vector. Changes to the expression levels of *Adcy5*, *Adcy8*, *Oprm1*, and other genes involved in the AC/cAMP pathway (e.g. PKA, Epac) were also analyzed, but no significant differences were seen for any of these genes in either tissue (Figure 2B-F).

**Figure 2.**
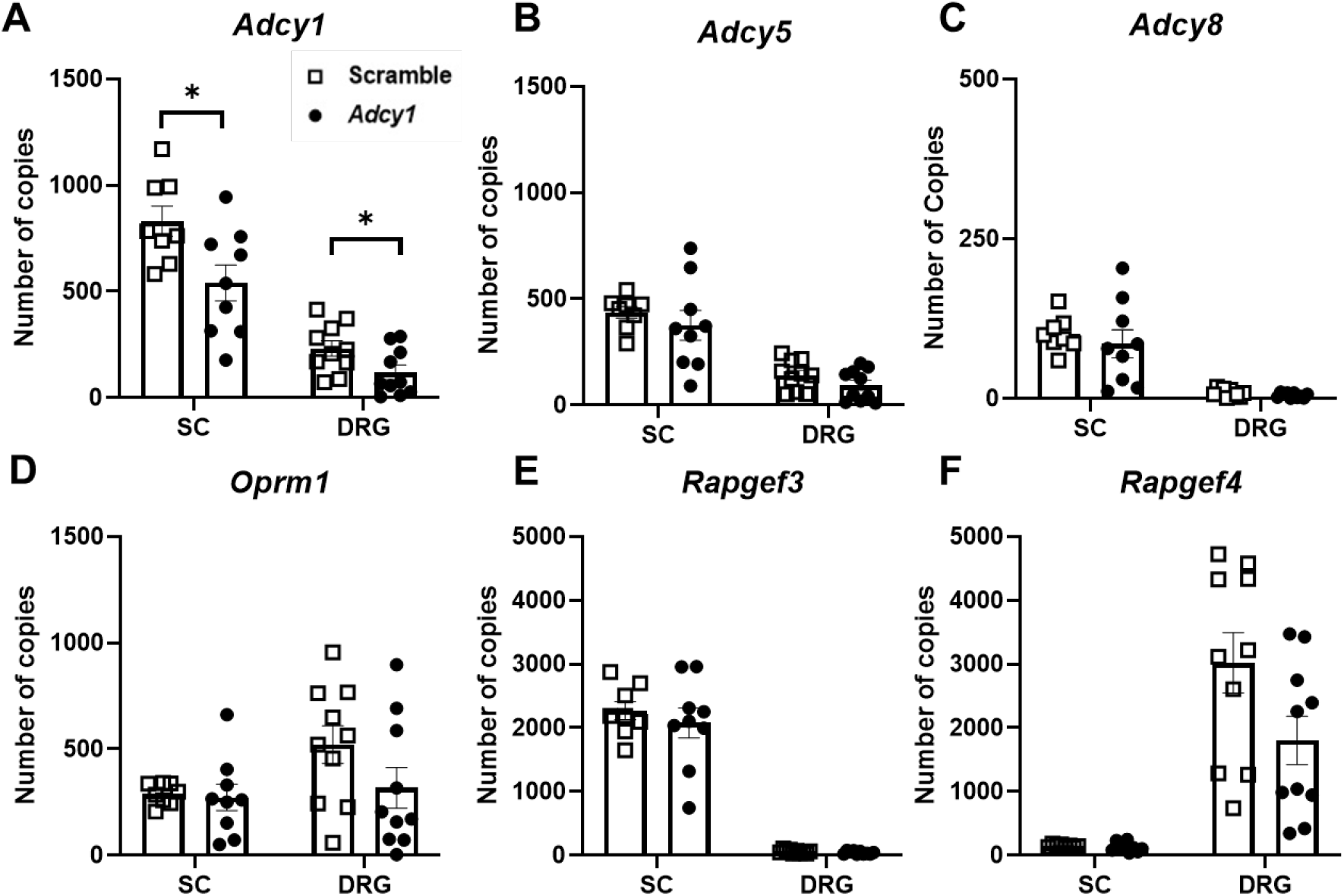
*Adcy1* expression is significantly decreased in the spinal cord and dorsal root ganglia after shRNA knockdown. Eight weeks after AAV9-GFP-U6-m-*Adcy1*-shRNA or AAV9-GFP-U6-scramble-shRNA intrathecal injection, qRT-PCR analysis was performed for *Adcy1* (A), *Adcy5* (B), *Adcy8* (C), *Oprm1* (D), *Rapgef3* (E), and *Rapgef4* (F) within the spinal cord (SC) and dorsal root ganglia (DRG). *Adcy1* was significantly decreased in AAV9-*Adcy1* (●) vector injected mice in both SC (A: unpaired t-test, p = 0.0201) and DRG (unpaired t-test, p = 0.0370) compared to AAV9-scramble (□) vector injected mice. Asterisk indicates statistical significance (p < 0.05). Data presented as mean ± SEM with an n=8-10/group.

Since continued agonist stimulation of the MOR increases AC1/cAMP activity, the *Adcy1* knockdown model was hypothesized to show an attenuation of morphine tolerance and opioid-induced hypersensitivity, but not necessarily acute morphine antinociception. An acute dose response curve indicated AAV9-*Adcy1* and AAV9-scramble treated mice had similar antinociceptive effects of morphine (Figure 3A; two-way ANOVA, F (1, 18) = 0.3231, p = 0.5768). This suggests that a single administration of morphine remains equally effective after knockdown of AC1. Five weeks post viral vector injections, mice were administered morphine twice daily in saline (15 mg/kg, sc) and MPW thresholds were measured twice daily, before the morning injections of morphine (Figure 3B) and 30 minutes post injection (Figure 3C). On Day 6, MPW thresholds were taken ~18 hours after last morphine injection (Figure 3B). AAV9-*Adcy1* injected mice had significantly higher MPW thresholds compared to AAV9-scramble injected mice both pre-morphine administration (Figure 3B; two-way ANOVA with Bonferroni’s *post hoc* test, F (1, 18) = 7.323, p = 0.0145, CI_Day6_ = −1.933 to −0.7565) and post-morphine (Figure 3C; two-way ANOVA with Bonferroni’s *post hoc* test, F (1, 18) = 5.847, p = 0.0264, CI_Day4_ = −2.034 to −0.1778, CI_Day5_ = −2.211 to −0.3548) indicating the knockdown of *Adcy1* not only attenuates the development of morphine tolerance but also the development of opioid-induced hypersensitivity.

**Figure 3.**
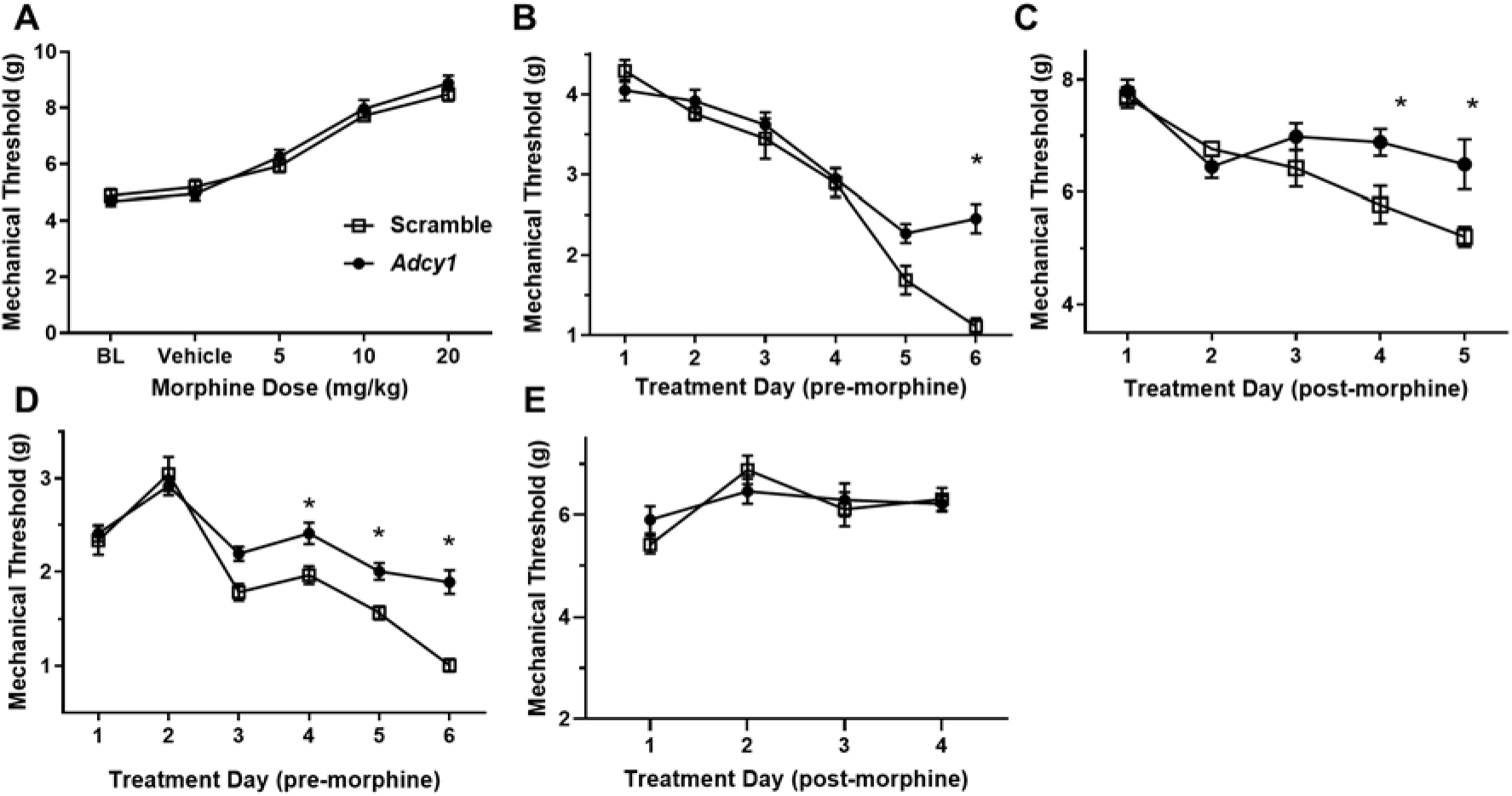
Intrathecal shRNA knockdown of *Adcy1* attenuates morphine tolerance and morphine induced hypersensitivity in mice. MPW thresholds were measured after intrathecal injection of either AAV9-GFP-U6-m-*Adcy1*-shRNA (●) or AAV9-GFP-U6-scramble-shRNA (□) four to six weeks post viral vector injections. During morphine dose response (A), baseline (BL) thresholds were measured before mice were given injections of vehicle. Increasing doses of morphine in saline were administered and MPW thresholds were measured 30 minutes post injection. No significant differences were seen between the two treatment groups. In a morphine tolerance paradigm, mice were injected twice daily with 15 mg/kg morphine in saline for five days (B-C) or were administered twice daily morphine in saline injections starting with 10 mg/kg and increasing by 10 mg/kg every day for four days until reaching 40 mg/kg on day four (D-E). Baseline measurements were taken every morning before and 30 minutes after morphine administration, with treatment day six (6) measurements taken ~18 hours after last morphine administration (B) and with MPW thresholds on days five and six measured ~18 hours and ~42 hours post morphine injection, respectfully (D). AAV9-*Adcy1* viral vector injected mice had significantly higher MPW thresholds during morphine tolerance testing compared to AAV9-scramble injected mice both pre-morphine (B; two-way ANOVA with Bonferroni’s *post hoc* test, F (1, 18) = 7.323, p = 0.0145, CI_Day6_ = −1.933 to −0.7565) and post-morphine (C; two-way ANOVA with Bonferroni’s *post hoc* test, F (1, 18) = 5.847, p = 0.0264, CI_Day4_ = −2.034 to - 0.1778, CI_Day5_ = −2.211 to −0.3548). AAV9-*Adcy1* viral vector injected mice had significantly higher MPW thresholds during baseline measurements of escalating morphine tolerance compared to AAV9-scramble injected mice pre-morphine (D; two-way ANOVA with Bonferroni’s *post hoc* test, F (1, 18) = 23.51, p = 0.0001, CI_Day4_ = −0.8649 to −0.03292, CI_Day5_ = −0.8556 to −0.02362, CI_Day6_ = −1.299 to −0.4668). (E) No significant difference in morphine efficacy is seen during escalating morphine tolerance. Asterisk indicates statistical significance (p < 0.05). Data presented as mean ± SEM with an n=10/group.

Using an escalating morphine tolerance model, mice were subjected to MPW latency testing while given twice daily injections of increasing doses of morphine in saline, starting with 10 mg/kg on Day 1 and increasing by 10 mg/kg daily until reaching 40 mg/kg on Day 4. The escalating morphine tolerance paradigm was used because the development of tolerance or opioid-induced hypersensitivity can lead to increased pain in clinic, with the usual consequence of escalating doses of opioids, either by prescription or self-medication[19]. MPW thresholds were measured every AM before (Figure 3D) and 30 minutes post-morphine administration (Figure 3E). On Day 5 and Day 6, MPW thresholds were measured ~18 hours and ~42 hours after the last morphine dose, respectfully (Figure 3D). AAV9-*Adcy1* injected mice exhibited significantly higher MPW thresholds than AAV9-scramble injected mice pre-morphine administration on Day 4, and on Days 5 and 6 when no morphine was administered (Figure 3D; two-way ANOVA with Bonferroni’s *post hoc* test, F (1, 18) = 23.51, p = 0.0001, CI_Day4_ = −0.8649 to −0.03292, CI_Day5_ = −0.8556 to −0.02362, CI_Day6_ = −1.299 to −0.4668). However, no significant difference is seen in MPW thresholds post-morphine administration between AAV9-*Adcy1* and AAV9-scramble injected mice (Figure 3E). This indicates that knockdown of *Adcy1* significantly attenuates opioid-induced hypersensitivity as well as opioid withdrawal in the escalating morphine tolerance model.

### *Intrathecal knockdown of* Adcy1 *improves mechanical hypersensitivity and non-evoked behaviors after CFA injection in mice*

Previous research has demonstrated that pharmacological inhibition of AC1 via ST034307 could provide analgesia in a mouse model of chronic inflammatory pain[4]. We devised a similar test for analgesic efficacy after *Adcy1-*shRNA treatment seven weeks after inoculation. One hind paw of each mouse was injected with CFA and MPW thresholds were measured three hours to one week after injection on both the injected (ipsilateral, Figure 4A) and non-injected (contralateral, Figure 4B) hind paws. AAV9-*Adcy1* injected mice had significantly higher MPW thresholds than AAV9-scramble treated mice on both the CFA injected paw (Figure 4A; repeated measures ANOVA with Bonferroni’s *post hoc* test, F (1, 18) = 6.157, p = 0.0232, CI_168hrs_ = −0.8329 to −0.08993) and the uninjected (right) hind paw (Figure 4B; repeated measures ANOVA with Bonferroni’s *post hoc* test, F (1, 18) = 9.148, p = 0.0073, CI_48hrs_ = −1.384 to −0.0513). This data indicates gene knockdown of *Adcy1* does provide some analgesic efficacy in the chronic inflammatory pain model 48 hours to one-week post-CFA administration.

**Figure 4.**
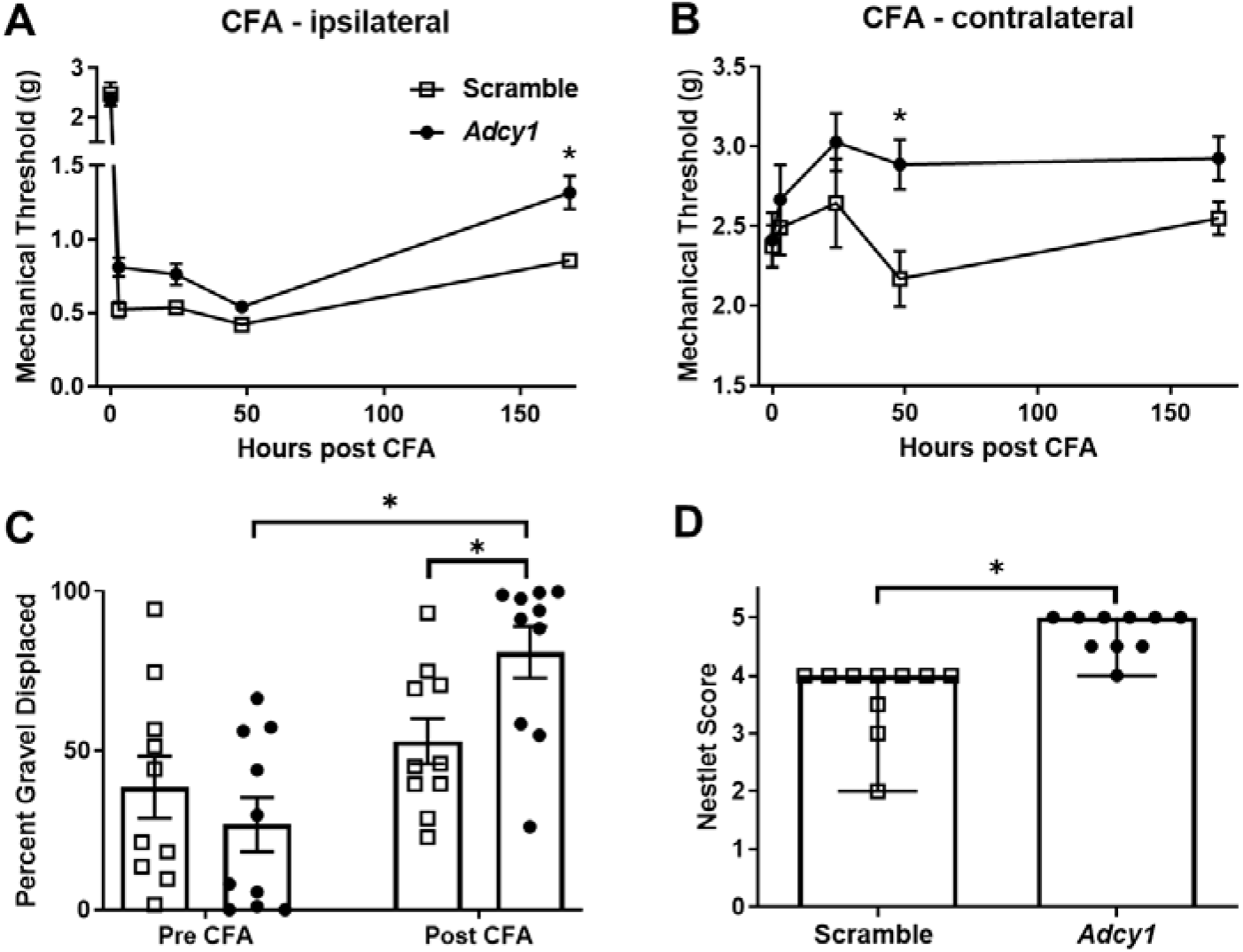
*Adcy1* intrathecal knockdown provides analgesia in mice with CFA induced inflammatory pain. Seven weeks post AAV9-GFP-U6-m-*Adcy1*-shRNA (●) or AAV9-GFP-U6-scramble-shRNA AAV9-scramble (□) injections, baseline MPW thresholds were measured before the left hind paw was injected with 20 μL undiluted CFA. MPW thresholds were then measured 3 hours, one day, two days, and one week post CFA injection on both the injected (ipsilateral) hind paw (A) and the uninjected (contralateral) hind paw (B). AAV9-*Adcy1* viral vector injected mice had significantly higher MPW thresholds than AAV9-scramble injected mice (□) on the injected paw (A; repeated measures ANOVA with Bonferroni’s *post hoc* test, F (1, 18) = 6.157, p = 0.0232, CI_168hrs_ = −0.8329 to −0.08993), and on the CFA uninjected paw (B; repeated measures ANOVA with Bonferroni’s *post hoc* test, F (1, 18) = 9.148, p = 0.0073, CI_48hrs_ = −1.384 to −0.0513). (C) AAV9-*Adcy1* and AAV9-scramble viral vector injected mice underwent burrowing testing three weeks post viral vector injections (Pre-CFA) and again seven weeks post viral vector injection (Post-CFA). AAV9-*Adcy1* viral vector injected mice had significantly higher percent gravel displaced during burrowing testing post-CFA than AAV9-scramble injected mice (C; two way ANOVA with Bonferroni’s *post hoc* test; F (1, 18) = 16.66; p = 0.0007; CI_Pre-CFA_ = −16.16 to 39.56, CI_Post CFA_ = −55.73 to −0.01139) and also significantly higher percent gravel displaced than their pre-CFA activity (two-way ANOVA with Bonferroni’s *post hoc* test; F (1, 18) = 16.66; p = 0.0007; CI_*Adcy1*_ = −83.03 to −25.03). (D) Nesting behavioral measures began seven weeks post viral vector injections corresponding to 3 days post CFA injection. AAV9-*Adcy1* viral vector injected mice had significantly higher nesting scores compared to AAV9-scramble injected mice (Mann Whitney U test; p < 0.0001). Asterisk indicates statistical significance (p < 0.05). Data in (D) presented as Median with Range, all others presented as mean ± SEM with n=10/group).

During this stage of chronic inflammation, non-evoked measures of pain and animal well-being including burrowing and nesting were examined to gain a better understanding of the impact of AC1 expression on behavioral measures during chronic inflammation. Both AAV9-*Adcy1* knockdown and AAV9-scramble treatment groups were subjected to burrowing testing three weeks and seven weeks post viral vector injections. No significant differences in burrowing behaviors were seen between the two treatment groups at the three weeks post viral vector injections (Figure 4C, Pre-CFA) but a significant difference was seen between the two treatment groups four days after CFA injection (Figure 4C, Post-CFA, two way ANOVA with Bonferroni’s *post hoc* test; F (1, 18) = 16.66; p = 0.0007; CI_Pre-CFA_ = −16.16 to 39.56, CI_Post CFA_ = −55.73 to −0.01139). A significant difference was also seen between the pre-CFA and post-CFA burrowing results for the AAV9-*Adcy1* viral vector treated group (Figure 4C, two-way ANOVA with Bonferroni’s *post hoc* test; F (1, 18) = 16.66; p = 0.0007; CI_*Adcy1*_ = −83.03 to −25.03).

Nesting behaviors were also conducted seven weeks post viral vector injections. A significant difference in nesting scores was seen between AAV9-*Adcy1* and AAV9-scramble injected mice (Figure 4D, Mann Whitney U test; p < 0.0001). Altogether, this data indicates the level of ongoing pain or discomfort may be decreased after AC1 knockdown and the loss of AC1 signaling may contribute greater functional motility during chronic pain.

### Knockdown of Adcy1 does not alter mobility or thermal nociception in mice

Additional behavioral tests were performed in order to see if a reduction of AC1 expression would affect other animal behaviors. Mice were subjected to both rotarod and open field assessments and thermal paw withdrawal testing three and four weeks post viral-vector injections, respectively. For rotarod testing, the total time on rotarod (Figure 5A) and maximum speed reached (Figure 5B) were not significantly different between AAV9-scramble and AAV9-*Adcy1* mice. No significant difference was seen between AAV9-scramble and AAV9-*Adcy1* mice during TPW testing (Figure 5C). In open field tests, no significant difference was seen between AAV9-Scramble and AAV9-*Adcy1* viral vector injected mice in distance traveled (Figure 5C), velocity (Figure 5F) and change in orientation angle (Figure 5G;). However, a small yet significant difference was seen in time spent immobile (Figure 5E; unpaired t-test, p = 0.0433) indicating that AAV9-*Adcy1* viral vector injected mice spent less time stationary compared to AAV9-scramble injected mice. Altogether, this data indicates the AAV9-*Adcy1* shRNA does not cause any major mobility changes in mice.

**Figure 5.**
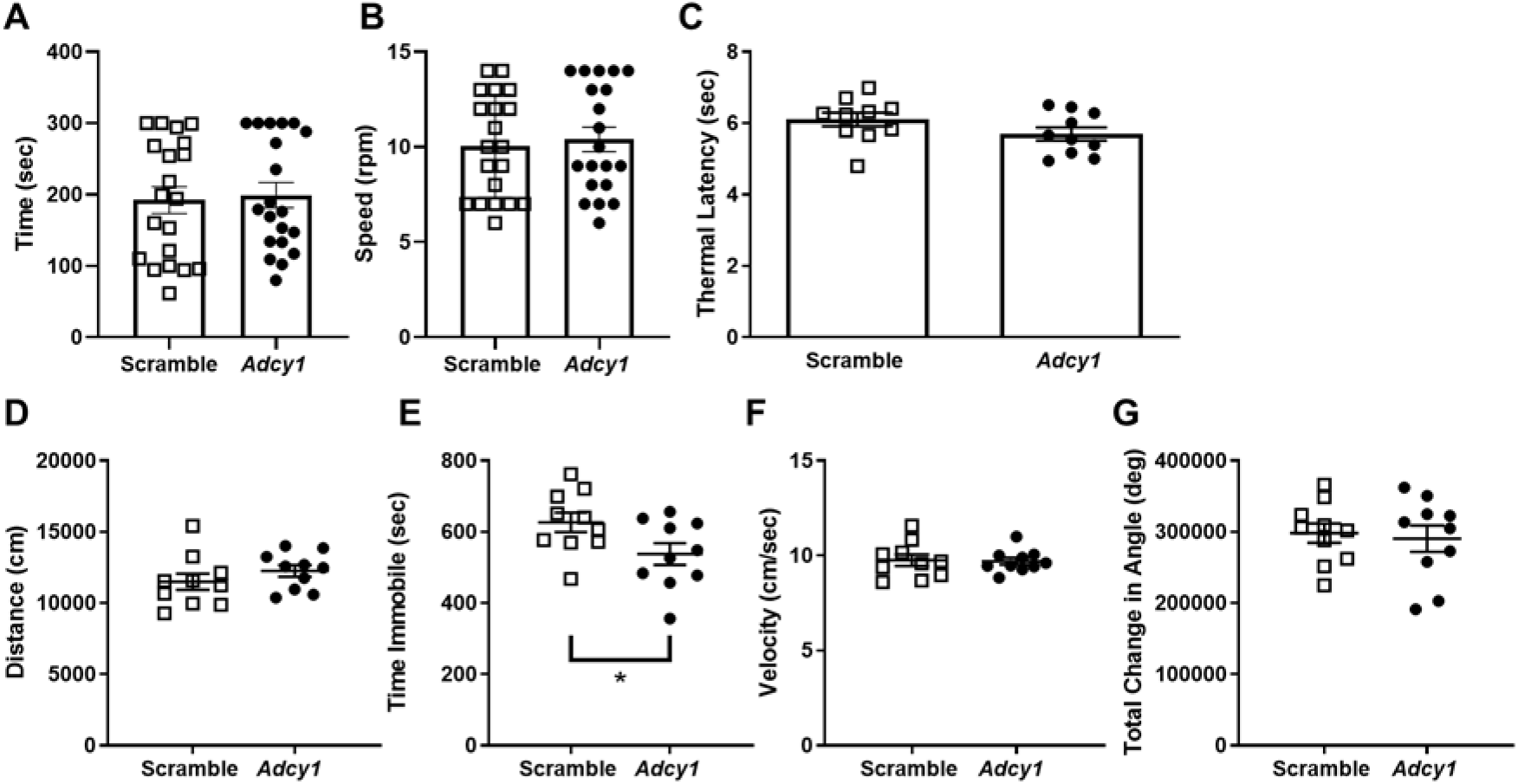
Animal mobility, thermal sensitivity and open field behaviors are not affected in mice with intrathecal knockdown of *Adcy1*. Three to four weeks post AAV9-GFP-U6-m-*Adcy1*-shRNA (●) or AAV9-GFP-U6-scramble-shRNA (□) intrathecal injections, mice underwent behavioral assessments to gauge mobility and the presence of deficits prior to morphine testing. For rotarod testing (A, B), mice were placed on the rotarod with an initial speed of 4 rpm increasing 1 rpm every 30 seconds to a maximum of 14 rpm. The maximum time (A) and revolutions per minute (B) were not significantly different across treatment groups. (C) For thermal paw withdrawal latency testing no significant difference is seen between AAV9-scramble and AAV9-*Adcy1* mice. For open field assessment (D-G), mice were placed inside arena and activity was recorded for 30 minutes. Distance traveled in cm (D), time spent immobile in seconds (unpaired t-test, p = 0.0433) (E), velocity in centimeters per second (F) and change in orientation angle in degrees (G) were all calculated for both AAV9-scramble and AAV9-*Adcy1* vector injected mice. No significant differences were seen between the two treatment groups in distance traveled, change in orientation angle and velocity. Data presented as mean ± SEM with an n=10/group.

### *AAV9-*Adcy1 *knockdown does not alter sciatic nerve conduction*

Viral inoculation was confirmed by fluorescence microscopy and the presence of GFP (Figure 6A). GFP signal was visualized in the sciatic nerves of all inoculated mice. Both right and left sciatic nerves were isolated and C-fiber compound action potentials (CAPs) were performed eight weeks post viral vector injection and CAPs were recorded. The downregulation of *Adcy1* did not have any impact on thresholds, amplitude, or conduction velocity of CAPs (Figure 6B-D.)

**Figure 6.**
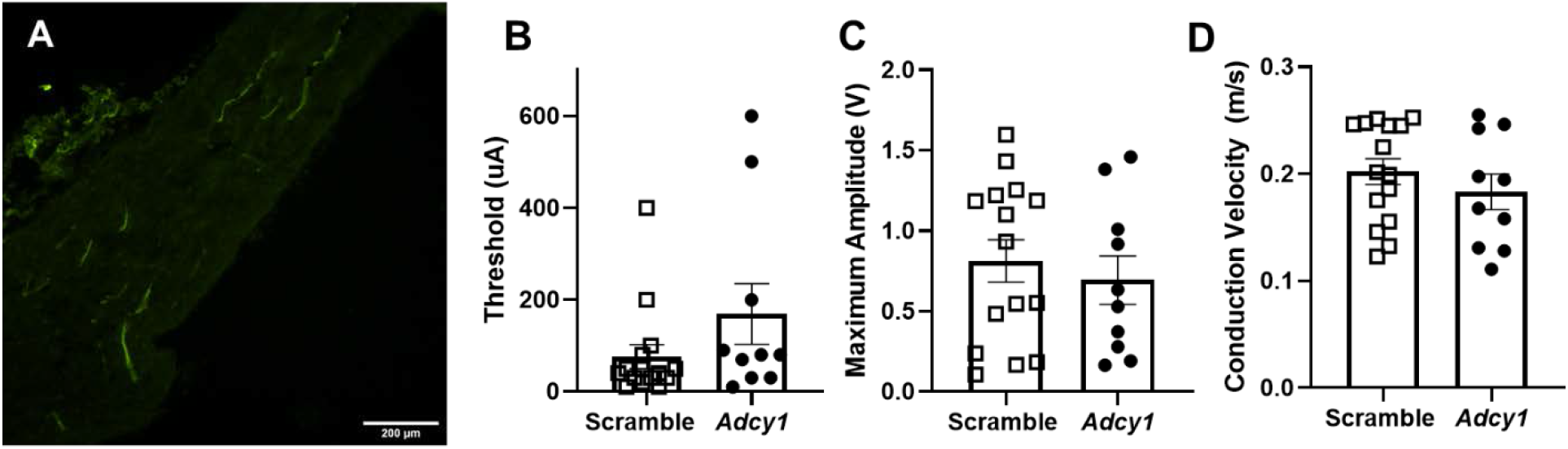
Knockdown of *Adcy1* does not impact sciatic nerve conduction. Eight weeks post intrathecal inoculation with either AAV9-GFP-U6-m-*Adcy1*-shRNA (●) or AAV9-GFP-U6-scramble-shRNA (□), sciatic nerves were removed and used for compound action potential recordings. (A) GFP signal visualized by fluorescence microscopy. Scale bar = 200 microns. The electrical thresholds (B), maximum CAP amplitude (C), and conduction velocity (D), were not significantly different between groups. Data presented as mean ± SEM with n=10-15/group.

## Discussion

The present study investigated the role of AC1 in a mouse model with regards to opioid tolerance, opioid-induced hyperalgesia, and inflammatory pain after CFA injection. Although all of the underlying mechanisms behind tolerance and opioid-induced hyperalgesia are not currently known, increased AC1 expression and activity has been suggested to be one of the major causative agents [8]. Our results indicate pharmaceutical inhibition of AC1 using ST034307 reduced opioid tolerance and attenuated morphine-induced hypersensitivity after increasing opioid administration. Intrathecal knockdown of *Adcy1* using a viral strategy was also effective at reducing morphine-induced hyperalgesia and withdrawal. The loss of *Adcy1* expression increased mechanical paw withdrawal thresholds, and improved burrowing and nesting behaviors after CFA intraplantar injection. This data suggests a reduction in the activity or function of AC1 may represent a novel analgesic target in addition to improving opioid withdrawal in patients taking opioids.

To date, there are nine membrane-bound AC isoforms, AC1-AC9 characterized in mammals and all nine have been confirmed in the nervous system [34]. AC1 is present in the brain, particularly in the cortex, hippocampus, and cerebellum, and historically has been thought to play a large role in learning and memory [12; 45]. AC1 is also present in the spinal cord [41] and in TrkA positive neurons in the DRG of mice [18]. A global loss in AC1 activity results in attenuated nocifensive behaviors after formalin hind paw injection and reduces pCREB activation in the superficial dorsal horn of the spinal cord [41]. The mechanisms that drive chronic pain are thought to be associated with opioid tolerance and both phenomena may arise from similar changes in intracellular signaling pathways in the peripheral and/or central nervous systems [20]. Along these lines, chronic morphine has been shown to produce a hypertrophied state of AC activity for AC 1, 6 and 8 *in vitro* [2]. The hypothesis that a selective AC1 inhibitor, ST034307, could also attenuate the development of morphine tolerance and hypersensitivity using an escalating dose paradigm was tested in our studies. Data presented here indicate that opioid-induced hyperalgesia was significantly attenuated after pharmacological inhibition of AC1, suggesting elevated activity of AC1 is responsible for the enhanced mechanical sensitivity upon cessation of daily morphine treatment.

Hypersensitivity and hyperalgesia seen in chronic pain and drug-induced hypersensitivity states most likely occur on multiple levels along sensory transmission pathways, from peripheral afferents, spinal cord synapses, and connectivity across midbrain and cortical cells. In chronic pain and opioid tolerant states, increased AC activity has been reported across the brain [28; 47] in addition to the spinal cord [42] and primary afferents [3; 46] which are thought to contribute to enhanced neurotransmission of nociceptive circuits. In order to determine if a localized decrease specifically targeted to AC1 activity in the spinal cord and primary afferent neurons could attenuate to opioid tolerance and inflammatory chronic pain, a genetic knockdown approach was used instead of a pharmacological one. Intrathecal delivery of AAV9 serotypes in live mice yield a high efficacy of transduction efficiency in DRG and lumbar spinal cord, while yielding sporadic labeling in the cortex and other peripheral tissues [32]. Static dosing of morphine (15 mg/kg, 2x daily, 5 days) and escalating doses of morphine over four days (10-40 mg/kg, 2x daily) both resulted in enhanced baseline mechanical sensitivity during the course of morphine administration. After intrathecal administration of AAV9-*Adcy1,* mice had higher mechanical paw withdrawal thresholds before (pre) and 30 minutes after (post) morphine administration. However, the acute morphine antinociception was not changed compared to control vector mice suggesting no change in acute pain responses which is similar to data obtained from AC1 knockout mice[41]. This important distinction between a lack of antinociception after acute morphine delivery and a significant enhancement of paw withdrawal thresholds after chronic morphine administration, indicate that adenylyl cyclase hypertrophy occurs after repeated stimulation of the MOR, and not after a single dose of an opioid, which has been a proposed paradigm for many years[43].

Systemic delivery of pharmacological inhibitors of AC1 have reduced hypersensitivity in neuropathic and inflammatory pain models in mice [4; 39]. The hypothesis that genetic knockdown of AC1 in the spinal cord and DRG could also attenuate inflammatory pain in mice was tested in our studies. Using a CFA model, AAV9-*Adcy1* mice had higher mechanical paw withdrawal thresholds compared to control mice seven days after CFA injection. This attenuation of mechanical hyperalgesia was also seen on the contralateral (uninjected) hind paws. During chronic pain states, it is possible that anatomical sites nearby also become sensitized to painful or non-painful stimulation as reported in previous rodent studies[6; 15]. The inhibition of AC1 appeared to attenuate mechanical hypersensitivity on either the ipsilateral or contralateral hindpaws, which indicate that pharmaceuticals targeting AC1 could also help attenuate pain sensitization beyond the primary zone of injury. The lack of analgesia seen during the initial phases after CFA administration (3-24 hrs), could be due to the role of adenylyl cyclases in enhanced transcription of pro-inflammatory molecules, which could take several days to manifest[35]. Alternatively, those results may be explained by the tissue specificity or overall level of the *Adcy1* knockdown when compared to the AC1 knockout mice[41].

Spontaneous pain and animal wellbeing after initiation of chronic pain is less frequently investigated than evoked measures, so our studies incorporated alternative testing measures. Studies have shown burrowing and nesting tests can be used to evaluate spontaneous pain or tonic pain in rodents [14; 30]. Data presented here indicate no appreciable differences between AAV9-*Adcy1* and AAV9-scramble mice in burrowing behaviors prior to CFA injection. However, four days after CFA induction, there is a significant improvement in burrowing behavior in AAV9-*Adcy1* treated mice. A similar difference was also seen in the nesting scores after CFA induction, with AAV9-*Adcy1* treated mice demonstrating significantly higher nesting scores than control mice. In previous studies, burrowing behavior is reduced in CFA inflammatory pain models in rats and can be reversed by ibuprofen[1]. Similarly, nesting behaviors are attenuated after CFA injection in mice which can be reversed by ketoprofen or low doses of morphine[31]. It is notable that no significant differences were detected between AAV9-*Adcy1* and AAV9-scramble mice in either the rotarod, thermal paw withdrawal latencies, or open field testing parameters, indicating intrathecal knockdown of AC1 does not affect acute thermal pain thresholds or affect general ambulatory behaviors. Transcriptional knockdown of AC1 in the sciatic nerves of mice was not measured, but significant changes in C-fiber compound action potential properties were not observed in this study. These data indicate behavioral changes seen in the AAV9-*Adcy1* animals may be restricted to the spinal cord and/or DRG, or loss of AC1 function does not impact axonal propagation of C-fiber action potentials.

In conclusion, small molecule inhibition of AC1 with ST034307 reduced morphine tolerance and hyperalgesia in mice. Similarly, knockdown of AC1 in the spinal cord and DRG reduced opioid-induced hypersensitivity after chronic administration of morphine in mice. Additionally, behavioral differences seen after *Adcy1* reduction appear specific to chronic pain-related behaviors, as animal locomotion and acute nociception were unaffected compared to controls. These studies suggest sensitized AC1 may represent a novel pharmaceutical target for the reduction of chronic pain and the attenuation of opioid-mediated adverse effects such as hyperalgesia. Further research into the intracellular targets of AC1 may provide new opportunities for new therapeutics in the future.

## Acknowledgements

Author contributions: Performed experiments and analyzed data: K.J, A.D. A.E. and A.H.K. Conceived experiments, designed and directed the studies: A.H.K. and V.J.W. Wrote the manuscript: K.J, A.D., V.J.W. and A.H.K.

This work was supported by grants from the National Institute of Drug Abuse (K01DA042902; A.H.K),and the Summer Undergraduate Research Program from the University of Minnesota (A.D.) and the TRIO McNair Scholars Program from the College of St. Scholastica (A.E.).

## Supplemental Files

**Supplemental Figure 1.**
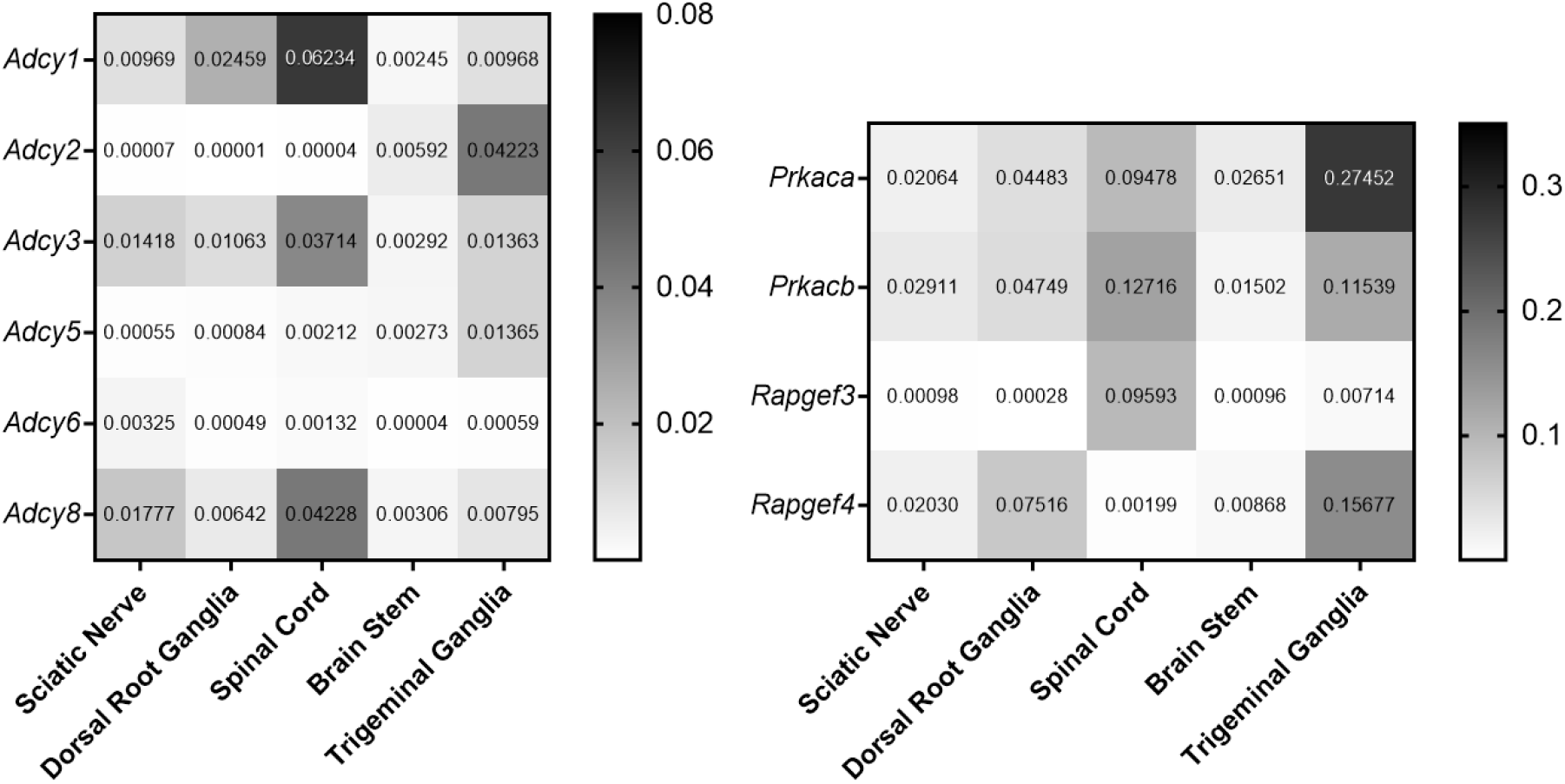
Heat map of genes expressed in saline-treated mice. (A) Adenylyl cyclases 1, 2, 3, 5, 6, and 8. (B) Protein kinase A catalytic (Prkaca) and regulatory subunits (Prkacb), Epac1 (Rapfgef3), and Epac 2 (Rapgef4). mRNA expression normalized to 18s expression in each tissue.

**Supplemental Table 1.**
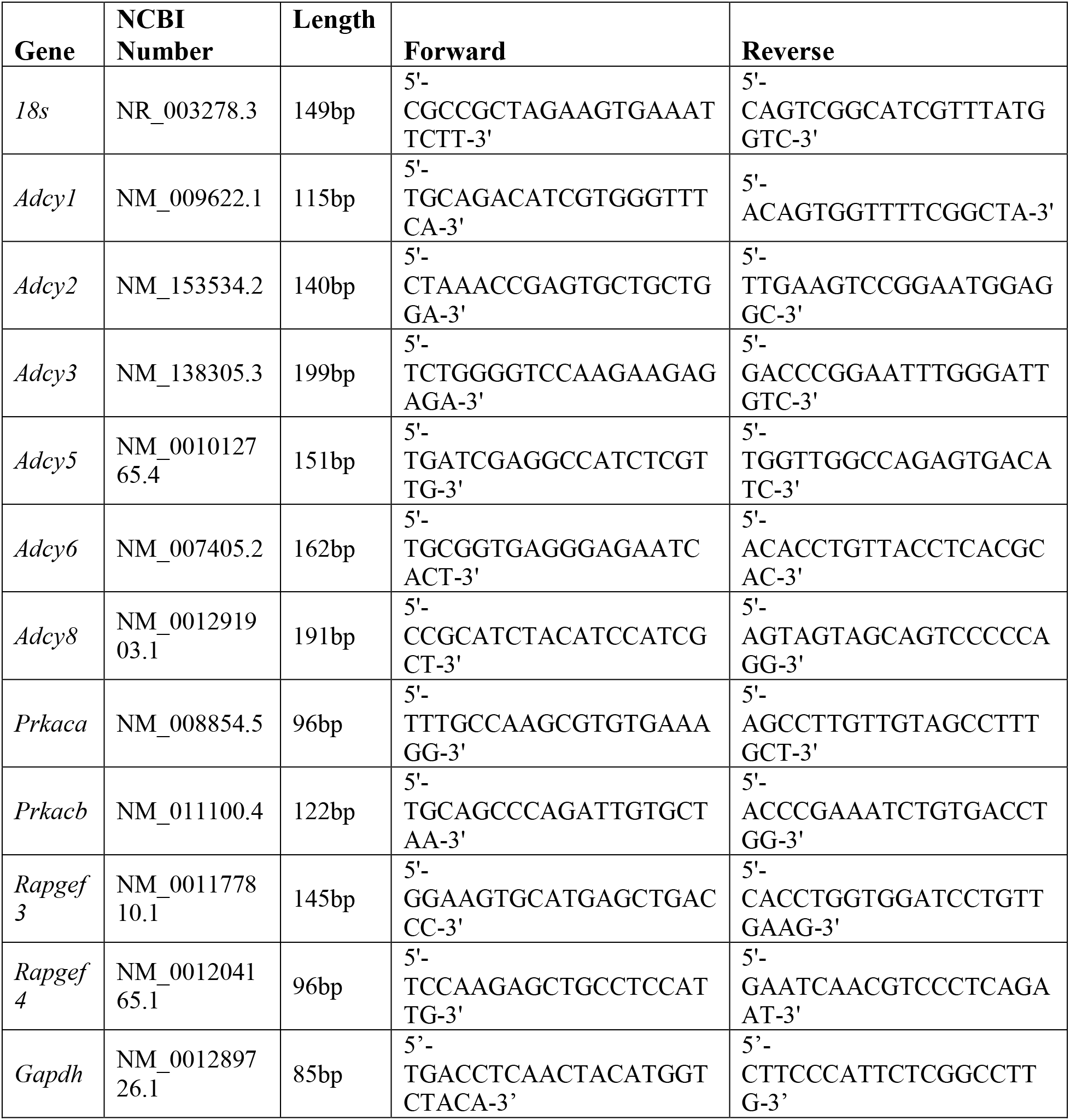
Gene Specific Primers Used for qRT-PCR. The NCBI gene accession number, resulting base pair length, and both the forward and reverse primers for each gene for qRT-PCR analysis.

